# Loss of α-actinin-3 confers protection from eccentric contraction damage in fast-twitch EDL muscles from aged *mdx* dystrophic mice by reducing pathological fibre branching

**DOI:** 10.1101/2021.10.01.462704

**Authors:** Leonit Kiriaev, Peter J. Houweling, Kathryn N. North, Stewart I. Head

**Affiliations:** School of Medicine, Western Sydney University, Sydney, NSW, Australia; Murdoch Children’s Research Institute, Melbourne, Victoria, Australia; Department of Paediatrics, University of Melbourne, Melbourne, Victoria, Australia

**Keywords:** *ACTN3*, *mdx*, EDL, DMD, branching

## Abstract

The common null polymorphism (R577X) in the *ACTN3* gene is present in over 1.5 billion people worldwide and results in the absence of the protein α-actinin-3 from the Z-discs of fast-twitch skeletal muscle fibres. We have previously reported that this polymorphism is a modifier of dystrophin deficient Duchenne Muscular Dystrophy. To investigate the mechanism underlying this we use a double knockout (dk)*Actn3KO*/*mdx* (dKO) mouse model which lacks both dystrophin and sarcomere α-actinin-3. We used dKO mice and *mdx* dystrophic mice at 12 months (aged) to investigate the correlation between morphological changes to the fast-twitch dKO EDL and the reduction in force deficit produced by an *in vitro* eccentric contraction protocol. In the aged dKO mouse we found a marked reduction in fibre branching complexity that correlated with protection from eccentric contraction induced force deficit. Complex branches in the aged dKO EDL fibres (28%) were substantially reduced compared to aged *mdx* EDL fibres (68%) and this correlates with a graded force loss over three eccentric contractions for dKO muscles (∼35% after first contraction, ∼66% overall) compared to an abrupt drop in *mdx* upon the first eccentric contraction (∼73% after first contraction, ∼89% after three contractions). In dKO protection from eccentric contraction damage was linked with a doubling of SERCA1 pump density the EDL. We propose that the increased oxidative metabolism of fast-twitch glycolytic fibres characteristic of the null polymorphism (R577X) and increase in SR Ca^2+^ pump proteins reduces muscle fibre branching and decreases susceptibility to eccentric injury in the dystrophinopathies.

## INTRODUCTION

Recent progress in gene-based therapies for Duchenne Muscular Dystrophy (DMD) show the potential to improve the outcomes for boys living with muscular dystrophy (Rodino-Klapac *et al*., 2013; Hoffman & McNally, 2014; Shieh, 2015; Bengtsson *et al*., 2016). However, clinical trials have yet to provide pathways for these genetic treatments to be translated into clinical practice (Aartsma-Rus & Muntoni, 2013; Fairclough *et al*., 2013; Hoffman & Connor, 2013; Lu *et al*., 2014). Some of the problems that occur are due to the genetic variability present among patients in the trials, thus there has been increased interest in identification of genetic modifiers to DMD. Studies have shown that the TGF-B pathway is a key activator of fibrosis in *mdx* mice and it is upregulated in DMD, where a late stage pathology of skeletal muscle is the presence of large amounts of non-contractile fibrotic material. The activity of the TGF-B pathway can be influenced by genetic modifiers such as osteopontin and LTBP4 (Ceco & McNally, 2013) potentially to reduce the amount of skeletal muscle fibrosis present in the dystrophinopathies. In a recent study by our group (Hogarth *et al*., 2017) investigating the common null polymorphism (R577X, rs1815739) in the *ACTN3* gene and its interaction with the mutated dystrophin gene in (DMD), we found young ambulant DMD boys had reduced muscle strength and slower 10 meter walking speeds when the gene was absent. Our study also found young double knockout (dk)*Actn3KO*/*mdx* (dKO) mice also had reduced muscle strength, but no differences in the degree of force deficit produced in fast-twitch muscles by eccentric contractions when compared to young *mdx*. In contrast, fast-twitch muscles from aged dKO mice were relatively protected from eccentric contraction force deficits compared to age matched *mdx* counterparts. Intriguingly, this suggests that the R577X polymorphism can have an age related ameliorating effect on muscle pathology in the dystrophinopathies. The polymorphism in the *ACTN3* gene occurs in an estimated ∼16% of people worldwide (North *et al*., 1999; Berman & North, 2010) resulting in an absence of the protein α-actinin-3 from the Z-discs of fast-twitch fibres. α-Actinin-3 is localised to the Z-discs of fast-twitch muscle fibres cross-linking the actin filaments of adjoining sarcomeres and interacting with a host of metabolic and signalling proteins (Lee *et al*., 2016). The main role of the Z-disc is to transmit movement and force generated by the contractile proteins, longitudinally to the tendons (Patel & Lieber, 1997). The absence of α-actinin-3 from fast-twitch fibres does not result in any muscle pathology, on the contrary it appears it may in fact be beneficial for endurance activities in humans (Yang *et al*., 2003), although it should be noted that several human studies (underpowered with interference from varied genetic backgrounds) do not provide support for the absence of α-actinin-3 improving human endurance (for a review see Houweling *et al*. (2018)). In order to circumvent this genetic variability in humans we developed an animal model, the *Actn3KO* mouse, which is genetically identical to its littermate controls apart from the absence of α-actinin-3. Using this model we have shown that α-actinin-3 significantly improves resistance to fatigue in fast-twitch muscles (Chan *et al*., 2008; Chan *et al*., 2011; Seto *et al*., 2011). We have also shown upregulated expression of the SERCA1 Ca^2+^ pumps in the fast-twitch muscles of *Actn3KO* mice that result in increased Ca^2+^ cycling at the intracellular Ca^2+^ store (the sarcoplasmic reticulum). This increased metabolic load required to pump the additional Ca^2+^ leak in α-actinin-3 deficient fast-twitch fibres may be a mechanism that explains the positive selection pressure during recent human evolution on the R577X polymorphism, as it would result in enhanced acclimatisation to the low environmental temperatures experienced by modern humans migrating out of Africa into the cold northern hemisphere over the last 60,000 years (Head *et al*., 2015). Further support of the positive evolutionary selection pressure in humans comes from our recent study showing that absence of α-actinin-3 is correlated to improved resistance to cold water immersion (Wyckelsma *et al*., 2021). In the *Actn3KO* mouse, there is no change in the fibre types found in fast-twitch muscles with respect to myosin heavy chain expression (Macarthur *et al*., 2008; Chan *et al*., 2011), however, fast fibres have reduced muscle fibre diameter (Macarthur *et al*., 2008) & mass (Chan *et al*., 2008; Chan *et al*., 2011), increased oxidative metabolism (Macarthur *et al*., 2007; Macarthur *et al*., 2008; Quinlan *et al*., 2010) and a slowed relaxation after contraction in young male animals (Chan *et al*., 2011). These skeletal muscle results can explain our recent findings (Hogarth *et al*., 2017) that in young ambulant boys with DMD and the *ACTN3* null polymorphism (R577X) is correlated with reduced muscle strength and longer 10m walk test time. Because of the implications for the presence of *ACTN3* in ∼16% of boys with DMD, in this current paper we verify and extended the animal experiments from Hogarth *et al*. (2017) in a new cohort of mice aged to 12 months. The 12 month age group was selected because we have previously demonstrated that by this age *mdx* fast-twitch EDL muscles are irreversibly damaged by a single eccentric contraction, and this irreversible eccentric force deficit is correlated with the number and complexity of branched dystrophic fibres present (Kiriaev *et al*., 2018; Kiriaev *et al*., 2021b). We found convincing correlative evidence that the age related protective effect conferred by the absence of α-actinin-3 is due to the reduction in the number and complexity of pathologically branched dystrophic fibres. Additionally, we propose that the mechanism by which the polymorphism R577X induces this protective effect in the dystrophinopathies is due to a switch in the fast-twitch fibres to a more glycolytic metabolism and an increase in the SERCA1 pumps, both of which are known to be protective in the dystrophinopathies (Goonasekera *et al*., 2011; Head *et al*., 2015; Mázala *et al*., 2015; Garton *et al*., 2018; Kiriaev *et al*., 2021a; Nogami *et al*., 2021).

## MATERIALS AND METHODS

### Ethics approval

Research and animal care procedures were approved by the Western Sydney University Animal Ethics Committee (A12346) and Murdoch Children’s Research Institute (MCRI) animal ethics committees and performed in accordance with the Australian Code of Practice for the Care and Use of Animals for Scientific Purposes as laid out by the National Health and Medical Research Council of Australia.

### Animals

*Mdx* (C57 BL/10SnSn-Dmd*mdx*/J) mice were obtained from Jackson Laboratories (Bar harbor, Me, USA). The *Actn3* knockout line created in this laboratory (Macarthur *et al*., 2007) was backcrossed onto the C57BL/10 background for 10 generations before being crossed with the *mdx* line to produce mice lacking both α-actinin-3 and dystrophin, (dk)*Actn3KO*/*mdx* (dKO). Animals were housed at a maximum of four to a cage in an environment controlled room with a 12h-12h light-dark cycle and had access to food and water ad libitum according to animal holding facility arrangements. Experiments were performed on 12 month old male mice with the dKO backgrounds (12.3 ± 0.09 months) and age matched *mdx* control mice (12.1 ± 0.1 months), genetic backgrounds were distinguished by genotyping. A total of 13 male mice were used in this study (n=5 aged *mdx* and 8 dKO mice).

### Muscle preparation

Animals were overdosed with isoflurane in an induction chamber, delivered at 4% in oxygen from a precision vaporizer. Animals were removed when they were not breathing and a cervical dislocation was immediately carried out. The fast-twitch EDL muscle was dissected from the hind limb in solution and tied by its tendons from one end to a dual force transducer/linear tissue puller (300 Muscle Lever; Aurora Scientific Instruments, ON, Canada) and secured to a base at the other end using 6-0 silk sutures (Pearsalls Ltd, Somerset, UK). The muscle was kept at room temperature during preparation in an organ bath containing Krebs solution with composition (in mM): 4.75 KCl, 118 NaCl, 1.18 KH_2_PO_4_, 1.18 MgSO_4_, 24.8 NaHCO_3_, 2.5 CaCl_2_ and 10 glucose, 0.1% fetal calf serum and bubbled continuously with carbogen (95% O_2_, 5% CO_2_) to maintain pH at 7.4. Krebs solution was used for dissecting, storage and contractile procedures.

The muscle was stimulated by delivering a current between two parallel platinum electrodes, using an electrical stimulator (701C stimulator; Aurora Scientific Instruments). All contractile procedures were designed, measured, and analysed using the 615A Dynamic Muscle Control and Analysis software (DMC version 5.417 and DMA version 5.201; Aurora Scientific Instruments). At the start of the experiment, the muscle was set to optimal length (L_o_), which will produce maximal twitch force, and measured using vernier muscle callipers. These contractile experiments were conducted on the EDL muscle and at a room temperature of ∼20–22°C.

### Initial maximum force

An initial supramaximal stimulus was given at 1ms, 125Hz for 1s, and force produced for muscles recorded as P_o_, the maximum force output of the muscle at optimal/resting length L_o_. Muscles then underwent an initial force frequency protocol, left to rest for 5 minutes (during which muscle length was reset to L_o_ with no stimulation in the organ bath), followed by the EC sequence and another 5 minute rest before a final force frequency protocol.

### Force frequency curve

Force-frequency curves were generated before and after the EC sequence to measure muscle contractile function throughout the study. Trains of stimuli were given 30s apart by different frequencies, including 2, 15, 25, 37.5, 50, 75, 100, 125, 150Hz at 1ms duration (width) width for 1s and the force produced was measured.

A sigmoid curve relating the muscle force (P) to the stimulation frequency (f) was fitted by linear regression to this data.

The curve had the equation:

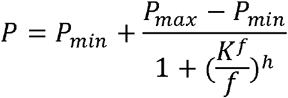

From the fitted parameters of the curve, the following contractile properties were obtained: force developed at minimal (P_min_) and maximal (P_max_) stimulation at the conclusion of the force-frequency curve. Half-frequency (K_f_) is the frequency at which the force developed is halfway between (P_min_) and (P_max_), and Hill coefficient (h) which provides a way to quantify the calcium binding affinity of the muscle during contraction. These were used for population statistics.

### Eccentric contractions

A series of eccentric (lengthening) contractions where the contracted muscle was stretched 10% from L_o_. At t=0s, the muscle was stimulated via supramaximal pulses of 1ms duration (width) and 125Hz frequency. At t=0.9s, after maximal isometric force was attained, each muscle was stretched 10% longer than their optimal length. The muscle was then held at this length for 2s before returning to optimal length. Electrical stimulus was stopped at t=5s. This EC procedure was performed 3 times at intervals of 3 minutes. The force measured at each EC was expressed as a percentage of the force produced during the first (initial) contraction.

### Twitch kinetics

Twitch force and kinetics were measured at two time points throughout the contractile protocol to compared differences between pre EC and post EC kinetics. The twitches were performed at 2Hz, for 1ms (width) and 1s duration immediately after initial maximum force (before) and 5minutes following the EC protocol (after). The following parameters were collected: peak twitch force expressed as a ratio of peak tetanic force, half relaxation time and time to peak.

### Work done

Work is an energy quantity given by force times distance, this measurement was used to provide a quantitative estimate of the EC-inducing forces. The work done to stretch the muscle was calculated from the force tracings through multiplying the area underneath the lengthening phase of the force tracing with the velocity of lengthening.

Brooks *et al*. (1995) examined the effect of various parameters on the force deficit produced by ECs, and found that the work done to stretch the muscle during the lengthening phase of an EC was the best predictor of the magnitude of the force deficit (Hunter & Faulkner, 1997; Stevens & Faulkner, 2000; Lindsay *et al*., 2020). Hence, this measure serves as a useful estimate of the mechanical stresses imposed on a muscle during an EC.

### Muscle mass

After contractile procedures were completed, the EDL muscle was removed from the organ bath and tendons trimmed. The muscles were blotted lightly (Whatmans filter paper DE81 grade) and weighed using an analytical balance (GR Series analytical electronic balance).

### Western Blot and Automated Western Blot protocol

Snap frozen extensor digitorum longus (EDL) muscles (n = 3 muscles / genotype) were lysed by sonication in 4% SDS lysis buffer containing 65mM Tris pH 6.8, 4% SDS, 1:500 Protease Inhibitor Cocktail solution (Sigma-Aldrich), 1X PhosSTOP Phosphatase Inhibitor Cocktail solution (Sigma-Aldrich) and Milli-Q water. Total protein concentrations were then assessed using Direct Detect (ThermoFisher Scientific) as per manufacturer’s instructions. Protein simple automated western blot was then performed using 0.2mg/mL of protein lysates, and primary antibodies including α-Sarcomeric actin (A2172, 5C5, Sigma-Aldrich, 1:10000) and SERCA1 (ab109899, Abcam, 1:5000), and their respective secondary antibodies in the 12-230 KDa capillary cartridge separation module (Protein Simple) as per manufacturer’s instructions. Protein analyses were performed using Compass software (Protein Simple). All results were then normalized to α-actin abundance and presented relative to healthy age matched WT controls (C57BL/10).

### Skeletal Muscle single fibre enzymatic isolation and morphology

Following contractile procedures and weighing, EDL muscles were digested in Krebs solution (without FCS) containing 3 mg/ml collagenase type IV A (Sigma Aldrich, MO, USA), gently bubbled with carbogen (95% O_2_, 5% CO_2_) and maintained at 37°C. After 25 minutes the muscle was removed from solution, rinsed in Krebs solution containing 0.1% fetal calf serum and placed in a relaxing solution with the following composition (mM): 117 K^+^, 36 Na^+^, 1 Mg^2+^, 60 Hepes, 8 ATP, 50 EGTA (note: internal solution due to chemically skinning by high EGTA concentration). Each muscle was then gently agitated using pipette suction, releasing individual fibres from the muscle mass and ensuring fibre distribution throughout the solution. Using a pipette 0.5ml of solution was drawn and placed on a glass slide for examination.

Where a long fibre covered several fields of the microscope view a series of overlapping photomicrographs were taken and these were then stitched together using the Corel draw graphic package. A total of 667 fibres from 9 EDL muscles were counted; 207 fibres from 4 aged *mdx* and 460 from 5 dKO animals. Only intact fibres with no evidence of digestion damage were selected for counting.

### Statistical analyses

Data was presented as means ± SD, Differences occurring between genotypes were assessed by individual student’s unpaired t-tests (unless specified) conducted at a significance level of 5%. All statistical tests and curve fitting were performed using a statistical software package Prism Version 7 (GraphPad, CA, USA).

## RESULTS

### Muscle mass, optimal length and twitch kinetics

The optimal muscle length L_o_ for both groups was identical, while dKO mass was slightly higher than aged *mdx* muscles (MD 2.466, 95% CI [-0.128, 5.059], P=0.06) this was not statistically different (Table 1). Before the EC protocol was administered the twitch kinetics of the aged *mdx* and dKO were the same, post EC the aged *mdx* showed a significant decrease in the time to peak and a depressed twitch/tetanus ratio c.f. the post EC dKO (see statistics in table 1). Interestingly, no differences in twitch half relaxation time was seen before and after the EC protocol between groups, however, overall there was an increase in both aged *mdx* (MD 4.6, 95% CI [1.274, 7.926], P=0.019, paired t-test) and dKO (MD 3.13, 95% CI [1.232, 5.018], P=0.006, paired t-test) half relaxation time post EC.

**Table 1.**
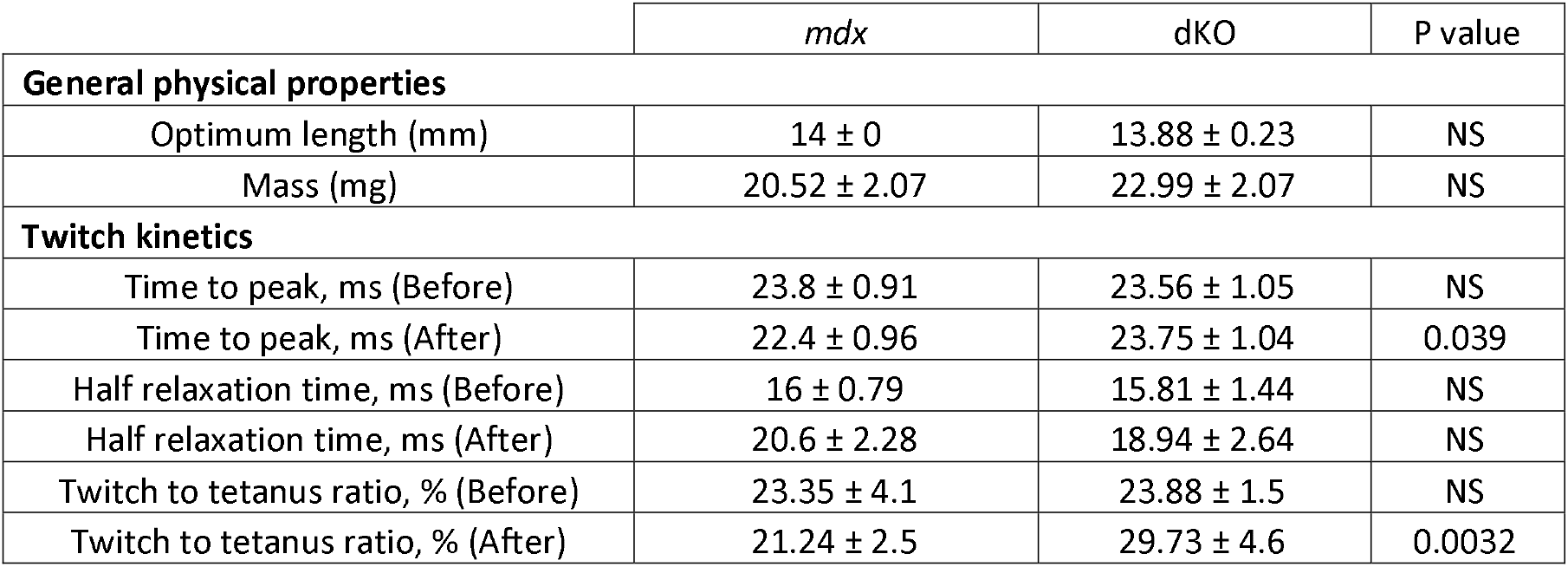
General physical properties and twitch kinetics of aged *mdx* and dKO EDL muscles. General physical properties: Muscle optimal length producing maximal force and muscle mass. Twitch kinetics: Values before and after EC are presented for Time to peak, Half relaxation time and twitch to tetanus ratio. All data shown are mean ± SD (n=5 Aged *mdx*; n=8 dKO). P-values shown are assessed by unpaired t-test with a significance established at 0.05, NS labelled for no statistical significance.

### Force-frequency parameters, half frequency and hill coefficient were unchanged by eccentric contraction protocol in aged mdx vers dKO

Figure 1A-C show aggregate force frequency curves generated from EDL muscle recordings at increasing frequencies from 2Hz-150Hz. The data from muscles were quantified by non-linear curve fitting using the sigmoidal equation given in the methods. For clarity, data for individual EDL muscles, were expressed as a percentage of maximum force and aggregated into single curves for each group. Fitted population parameters are derived from the curve fitting. Half frequency (Figure 1D) is the stimulation frequency at which the muscle develops force which is halfway between its minimum and maximum forces, the hill coefficient (Figure 1E) is a measure of the slope of the curve. Following EC significant changes occurred in the aged *mdx* EDL (but not dKO) with a ∼10% increase in the half frequency, resulting in a higher post EC half frequency between groups (MD 8.137, 95% CI [2.947, 13.33], P=0.0054) (Figure 1D), and a ∼48% decrease in the slope (hill coefficient) resulting in a lower post EC hill coefficient between groups (MD -3.015, 95% CI [-4.439, -1.591], P=0.0007)(Figure 1E). From the curve fitting we calculated a post EC force deficit of ∼67% for dKO c.f. ∼87% in aged *mdx* (MD -19.94, 95% CI [-3.685, -36.2], P=0.021) (Figure 1B & C).

**Figure 1.**
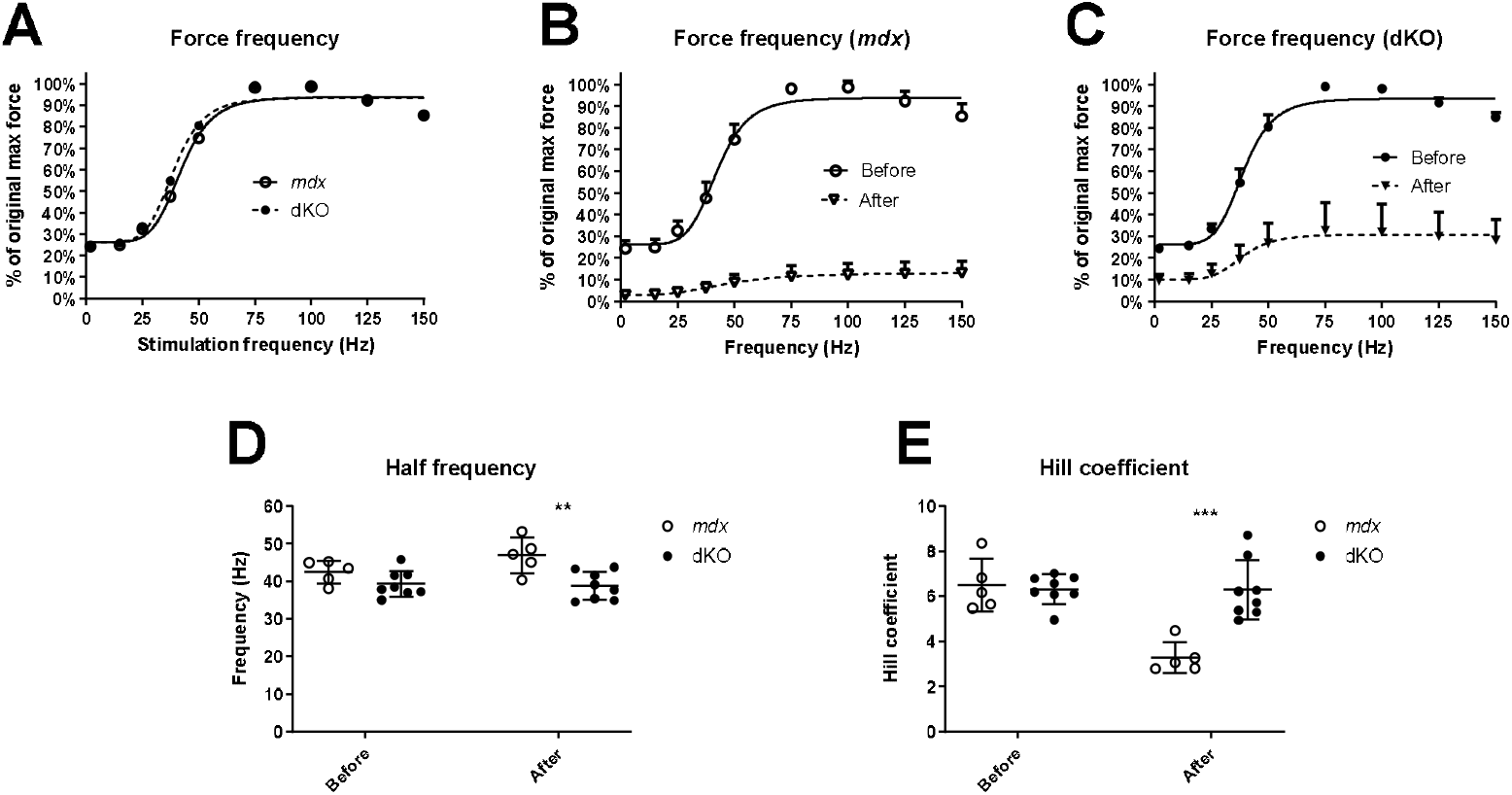
Force frequency curves and fitted parameters. A: Force frequency data from individual EDL muscles were aggregated to produce a single curve for aged *mdx* and dKO mice to visualize differences between genotypes prior to EC. SD omitted for clarity in A but provided later in B & C (top only). B and C: Force frequency curves obtained before (solid line) and after (dashed line) the EC protocol for aged *mdx* and dKO respectively. D: Scatterplots of Half frequency before and after EC for aged *mdx* and dKO. E: Scatterplots of Hill coefficient before and after EC for aged *mdx* and dKO. All data shown are mean ± SD (n=5 Aged *mdx*; n=8 dKO). P-values shown are assessed by unpaired t-test with a significance established at 0.05, ***0.0001<P<0.001 and **0.001<P<0.01.

### dKO EDL muscles are protected from eccentric contraction induced force deficits despite undergoing a greater “work done” during eccentric contractions

The isometric force deficits after each EC are expressed as a portion of starting force in Figure 2A. The EC force deficit occurred incrementally in the EDL muscles from the dKO with a ∼36% force deficit on the first EC, in contrast aged *mdx* EDL muscles showed an abrupt ∼75% force deficit on the first EC (MD -36.83, 95% CI [-51.82, - 21.85], P=0.0002) (Figure 2B left). After the three eccentric contractions EDL muscles from dKO mice lost ∼66% of starting force c.f. ∼88.58% EC force deficit in aged *mdx* (MD -22.06, 95% CI [-34.25, -9.88], P=0.0021) (Figure 2B right). We calculated the work done as the area under the force curve during the active lengthening phase of the first EC. Figure 2C illustrates that the work done during the first EC was significantly greater for EDL muscles in dKO animals compared with age matched *mdx* controls (MD 122.54, 95% CI [48.77, 196.3], P=0.0038).

**Figure 2.**
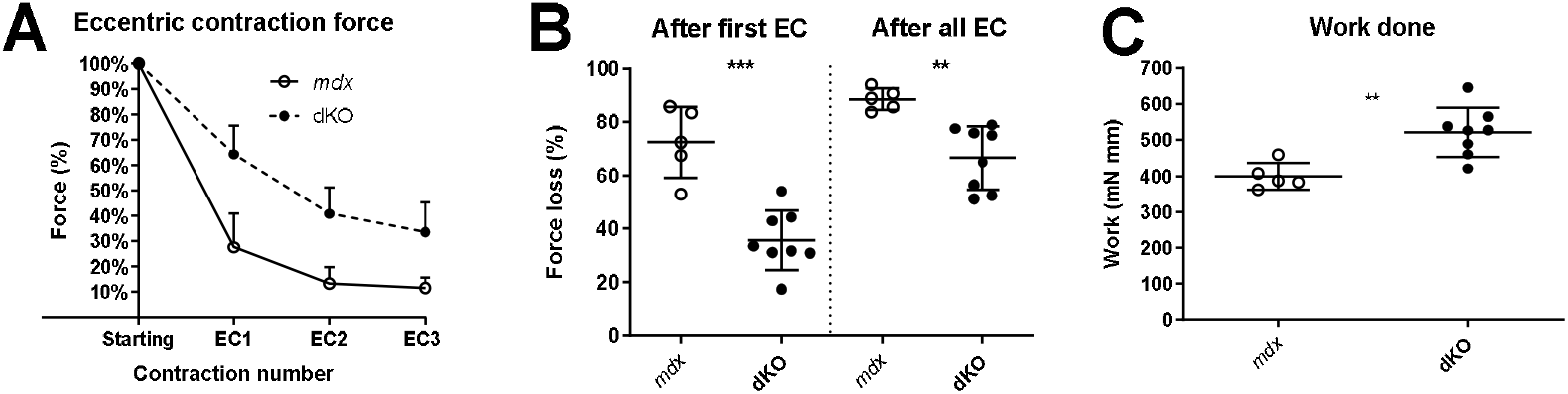
Force loss from EC and work done at 10% strain. A) XY plot of force loss across the 3 eccentric contractions expressed as a percentage of starting force for aged *mdx* and dKO EDL. Note: SD presented as top half only. B) For clarity scatterplots of force loss as a consequence of the first 10% EC (left) and after all 3 eccentric contractions (right) have been provided. C) Scatterplot showing work done calculated during the first 10% EC for aged *mdx* and dKO EDL. All data shown are mean ± SD (n=5 Aged *mdx*; n=8 dKO). P-values shown are assessed by unpaired t-test with a significance established at 0.05, ***0.0001<P<0.001 and **0.001<P<0.01.

### There are less complex branched fibres in EDL muscles from dKO mice which correlates with the protection from eccentric contraction force deficits

The number and complexity of fibre branching found in EDL muscles of dKO and age matched *mdx* control groups are shown in Figure 3 (note these counts represent all fibres counted and there are no error bars). In the aged *mdx* EDL around 97% of fibres contain branches; ∼6% of fibres contain a single branch, ∼23% contain 2 or 3 branches and ∼68% of all fibres develop complex branches (4 or more branches per fibre). In comparison both the number of branched fibres and the complexity of branching is reduced in dKO EDL. Approximately 89% of dKO muscle fibres contain branches; ∼22% of fibres contain a single branch, ∼39% contain 2 or 3 branches and ∼28% of all fibres develop complex branches (4 or more branches per fibre).

**Figure 3.**
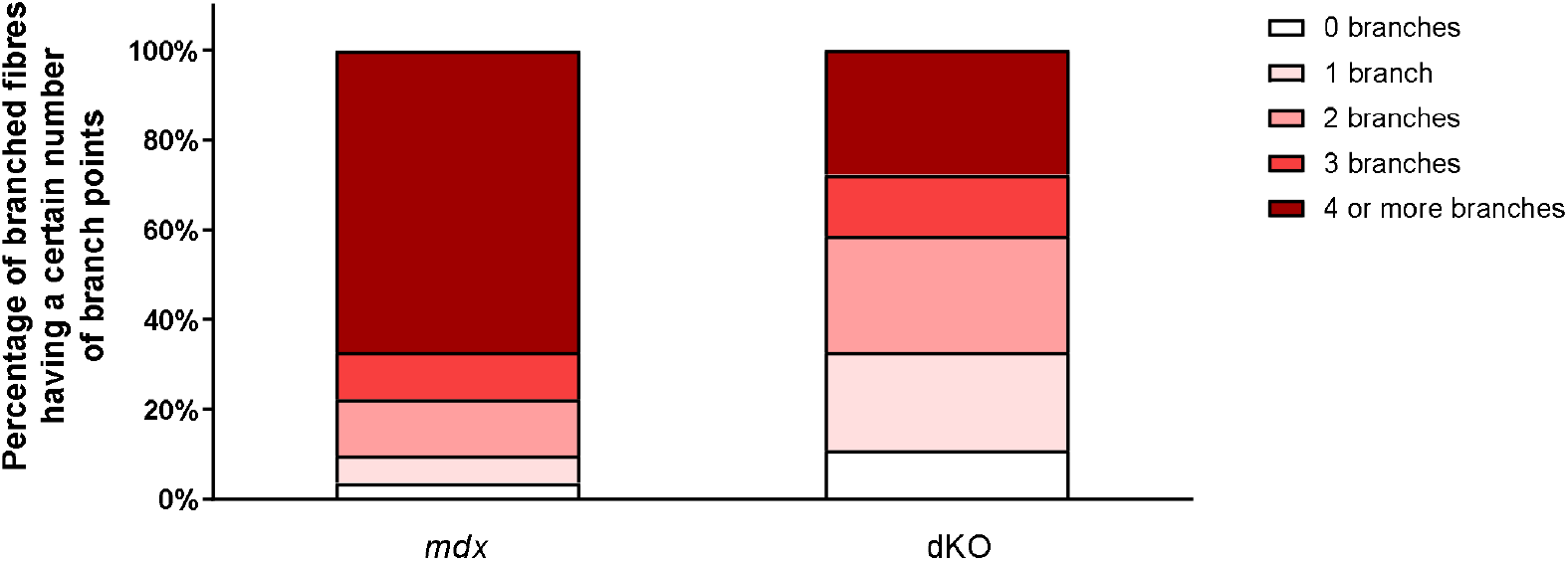
The degree of fibre branching in EDL muscles as a percentage of all fibres by categorizing them based on number of branch points. Note this is a bar graph of all fibres counted and as such is an absolute measure, all photomicrographs in Figures 4-6 were taken from these fibre counts. A total of 667 fibres from 9 EDL muscles were counted; 207 fibres from 4 aged *mdx* and 460 from 5 dKO animals. Only intact fibres with no evidence of digestion damage were selected for counting.

### Light microscope morphology of enzymatically isolated single fibres

Representative images of intact muscle fibres taken at magnification (X100) on a light microscope demonstrate the complexity of fibre branching in the aged *mdx* EDL (Figure 4). Panels A-B show some examples of simple branched fibres (1 or 2), panels C-E show more complicated fibre branches (3-4) and panels F-I are examples of long complex branched fibres (4+) with multiple offshoots along the length of the dystrophic fibre.

**Figure 4.**
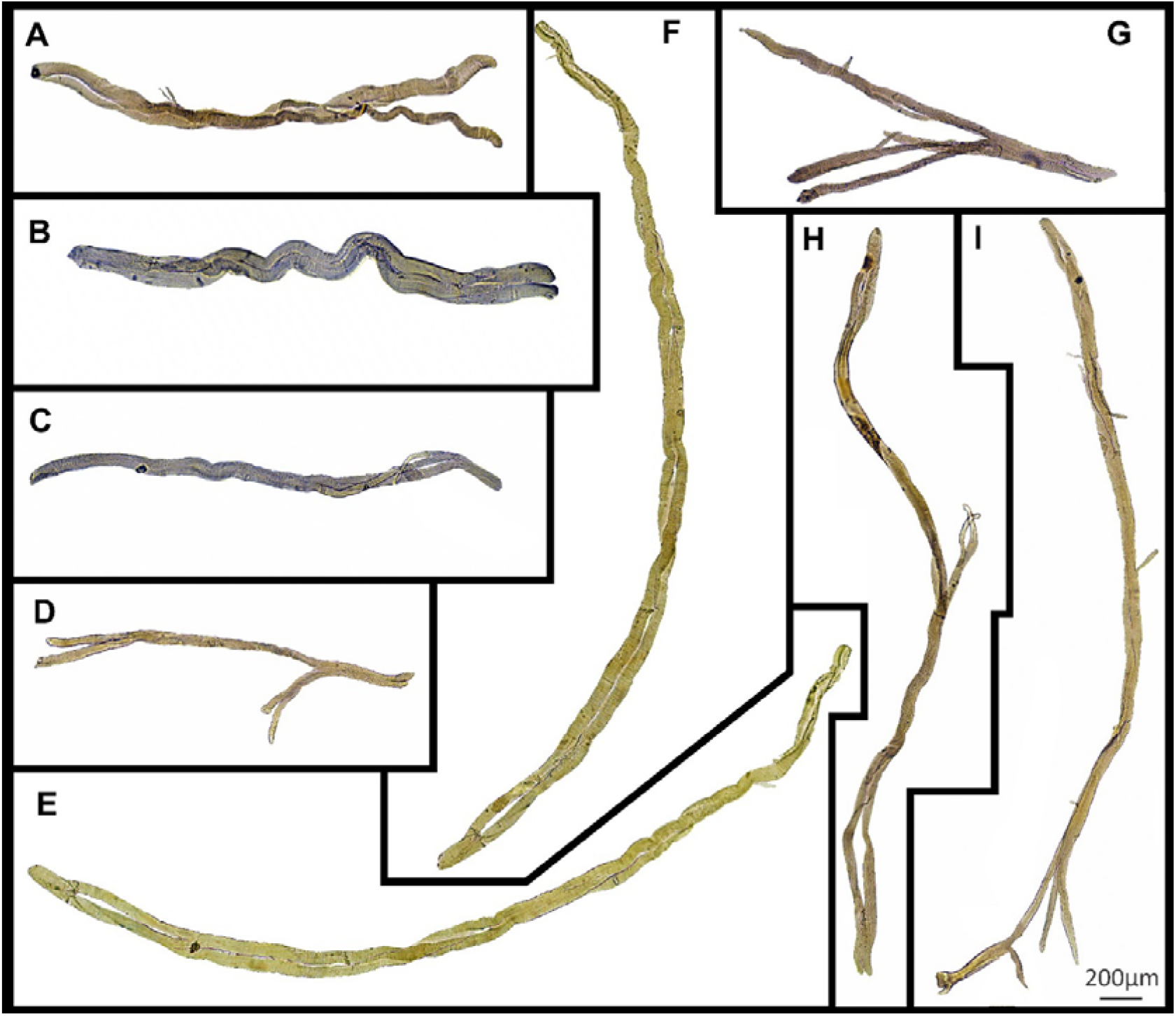
Examples of light microscope images (X100) of enzymatically digested EDL muscle fibres taken from aged *mdx* mice (n=2 muscles). Note: the fibres in Figures 4&5 have been stitched together at the same magnification from photomicrographs taken from overlapping fields of view to capture a large portion of the fibre, scale bar provided at 200µm in the bottom right corner applies to all panels. Background debris from the digest process have been cleared to focus on the muscle fibre. To see examples where debris have not been removed see Kiriaev et al. (2018). Simple branched fibres (1 or 2 branches) can be seen in panels A-B, more complicated fibre branches (3-4 branches) are seen in panels C-E and panels F-I show examples of long complex branched fibres (4+ branches) with multiple offshoots along the length of the fibre.

Images of single EDL fibres from dKO mice taken at magnification (X100) are shown in Figure 5. Figure 5 Panels A & D are examples of straight fibres with no branches and panels H & L are examples of more complex fibres found in the dKO muscle. Most dissociated fibres we observed resemble the morphology seen in all other panels in Figure 5, containing 1-3 branches per fibre. When comparing these light microscope images of dissociated fibres Figure 4 c.f. Figure 5, it’s evident that the complexity of fibre branching is reduced in dKO mice compared to aged *mdx* counterparts.

**Figure 5.**
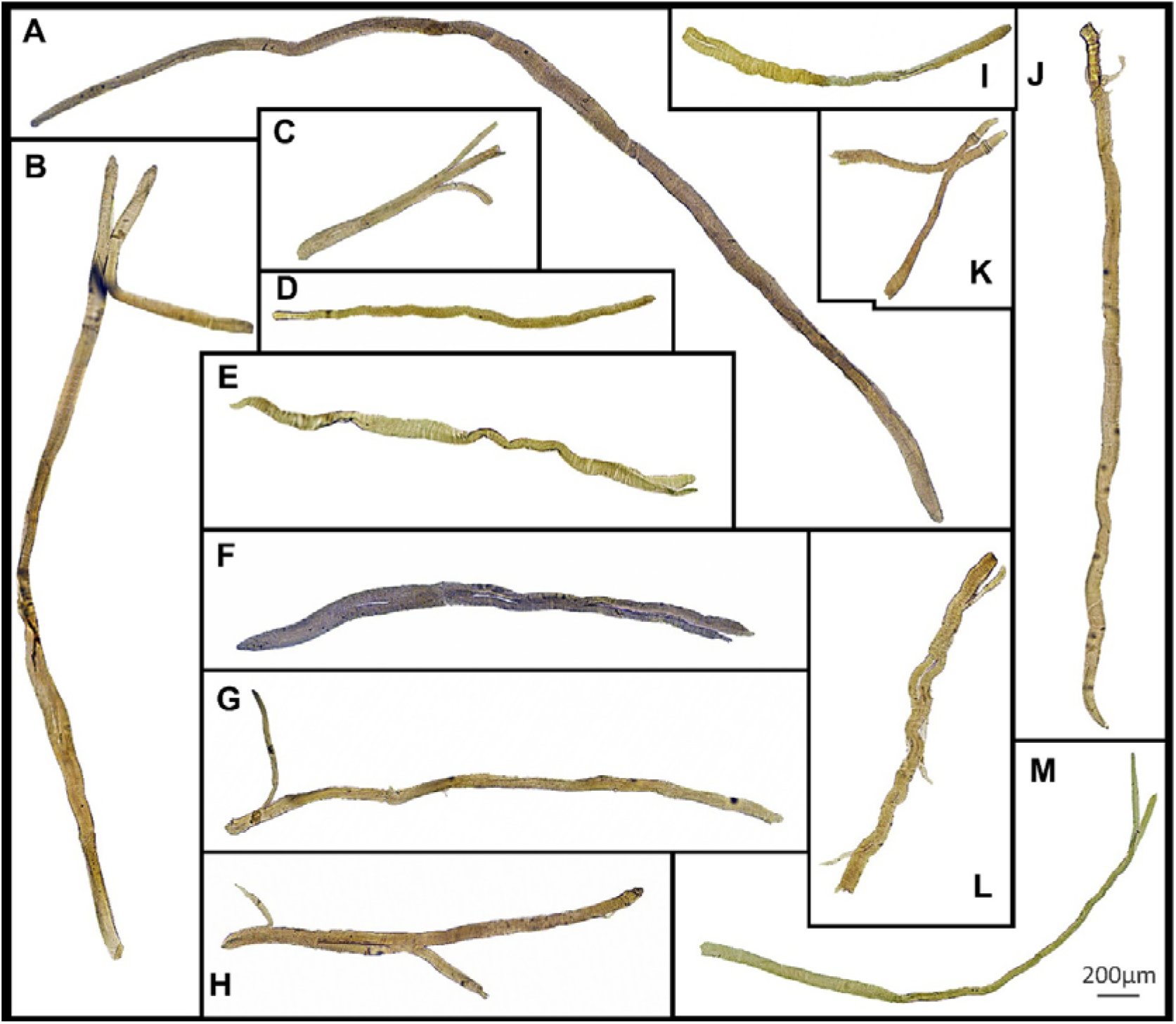
Examples of light microscope images (X100) of enzymatically digested EDL muscle fibres taken from dKO mice (n=3 muscles). Scale bar provided at 200µm in the bottom right corner applies to all panels. Straight fibres with no branching are seen in panels A&D as well as examples of complex branched fibres in panels H&L were also seen in the dKO EDL muscles. The majority of dissociated fibres resemble the morphology in all other panels containing between 1-3 branches per fibre.

Figure 6 shows some examples of magnification (X100) light microscope images from dKO EDL muscles where branch points can be seen at higher resolution and contrast. The branch pattern in these dKO fibres range from simple splits or offshoots towards the end of the fibre shown in panels A, G, I & J, to splits that rejoin mid fibre in panels B, C, F, H, K and L as well as tapered fibres in panels D & E. Cartoon inserts have been added to each image to help identify broken fibres (Red) and super-contracted areas (Yellow).

**Figure 6.**
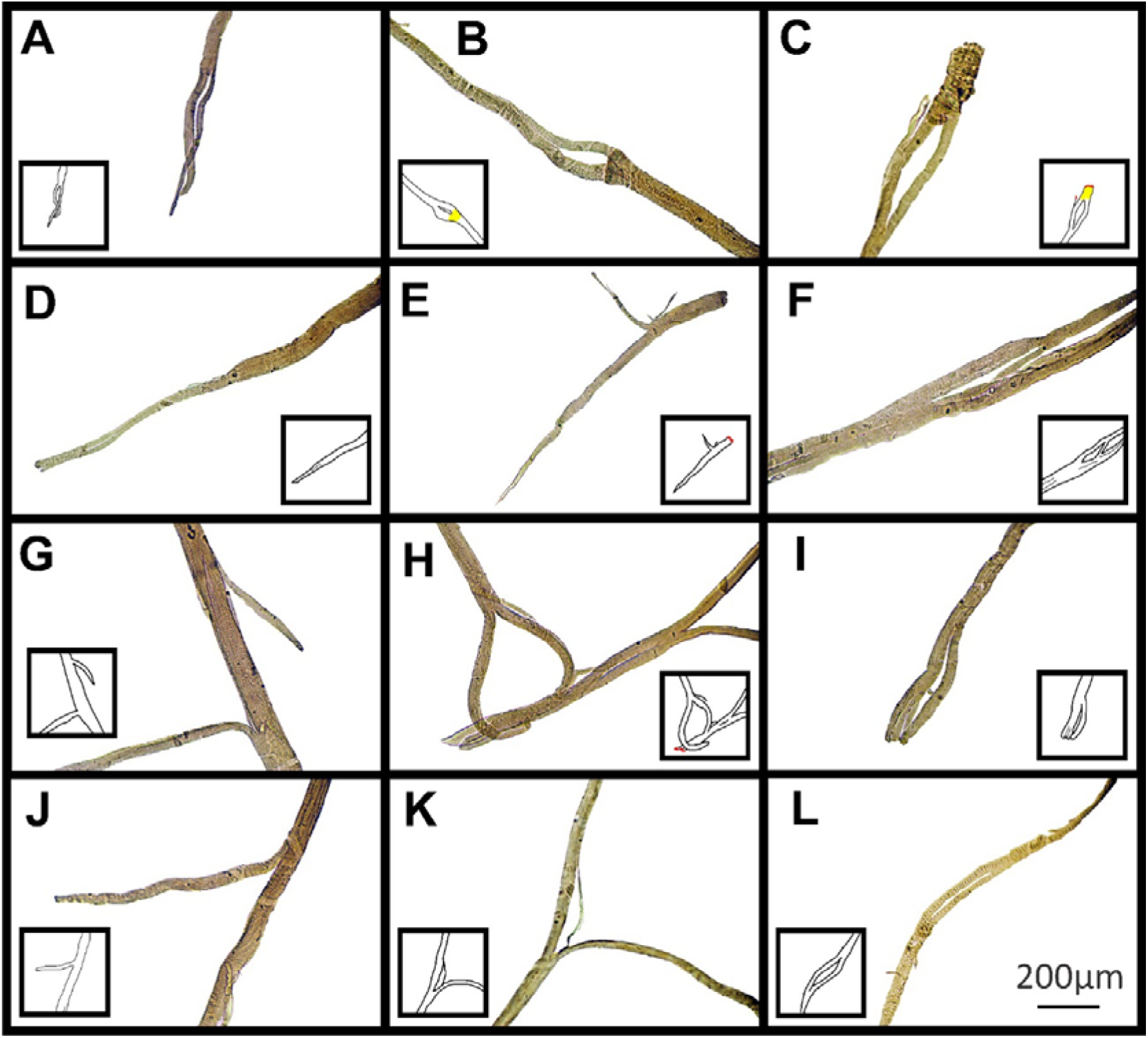
Examples of light microscope images (X100) focusing on the fibre branch point in dKO muscle fibres over one field of view. Cartoon inserts have been added to help visualize the various examples of branching in each image also highlighting broken (Red) and necrotic areas (Yellow). Branchpoint patterns such as simple splits or offshoots towards the end of the fibre shown in panels A, G, I & J to splits that re-join in the middle of fibres shown in panels B, C, F, H, K and L as well as tapered fibres in panels D & E. Scale bar provided at 200µm in panel L applies to all of figure 6.

### SERCA1 expression

We have previously reported that one consequence of *Actn3KO* in fast-twitch muscle is an increase in the expression of the fast SR Ca^2+^ pump SERCA 1. Using the Western Blot technique we analysed the level of SERCA1 expression in aged dKO mice and age matched *mdx* (Figure 7). We observed a 2 fold increase in the amount of SERCA1 protein in EDL muscles collected from dKO mice compared to age matched *mdx* (MD 1.033, 95% CI [0.736, 1.33], P=0.0006).

**Figure 7.**
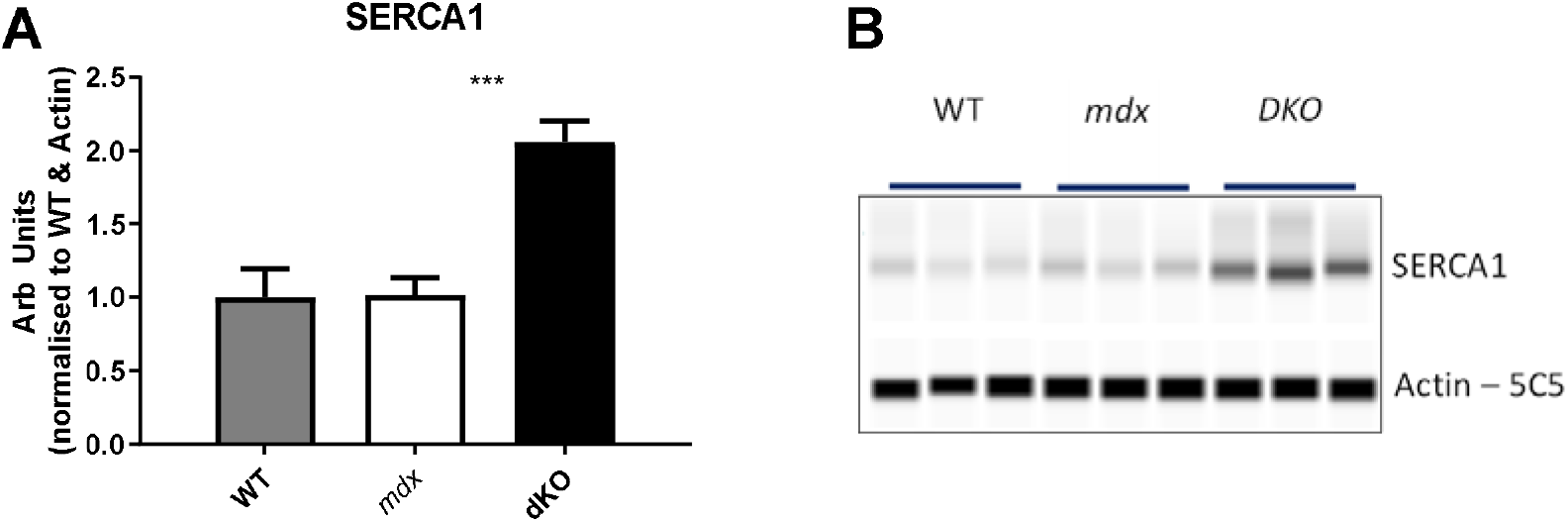
Expression of SERCA1 in age matched wild type, *mdx* and dKO EDL muscles. A: SERCA1 protein densitometry values normalized to α-skeletal actin and the average of all WT samples. B: Representative western blots that correspond to A. All data shown are mean ± SD (n=3 WT; n=3 Aged *mdx*; n=3 dKO). P-values shown are assessed by unpaired t-test with a significance established at 0.05, ***0.0001<P<0.001.

## DISCUSSION

### Muscle mass, optimal length and twitch kinetics

At base line these were the same in EDL muscles from 12 month old *mdx* and dKO mice. This supports our earlier reports that the *Actn3KO* polymorphism does not alter fibre type distribution or number of fibres in fast-twitch muscle at 12 months of age (Seto *et al*., 2011). Muscle twitch kinetics are an important parameter to measure as Peczkowski *et al*. (2020) and colleagues reported twitch kinetics yielded greater statistical significance compared to previously published maximal tetanic force measurements, and proposed the use of kinetics measurements to allow for use of smaller animal groups in *mdx* mouse studies. Thus, given the number of animals used in the present study we can be confident that the *Actn3* polymorphism in our dKO mice has not altered the contractile properties of the fast-twitch EDL muscle when compared to age matched *mdx* EDL muscles. However, in relation to kinetic parameters, as detailed in Figures 1 & 2, after the eccentric contraction protocol there are significant differences between the aged *mdx* and dKO mice. Twitch time to peak (MD 1.35, 95% CI [0.084, 2.616], P=0.039) and twitch/tetanus ratios (MD 8.45, 95% CI [3.51, 13.46], P=0.0032) were significantly higher in dKO mice (Table 1). Given it has been reported that a low twitch/tetanus ratio is characteristic of slow-twitch muscle fibres (Celichowski & Grottel, 1993) we can draw the conclusion that the *Actn3* polymorphism has been protective against EC induced force deficits in fast-twitch fibres from the dKO while in the age matched *mdx* the fast-fibres have been damaged by the EC protocol, leaving the slower fibres operational, explaining the slower twitch/tetanus ratio and time to peak (Table 1).

### Force-frequency parameters

The half frequency and hill coefficient are a measure of the sensitivity of the contractile proteins to Ca^2+^. The lower the half frequency and the higher the hill coefficient, the greater the degree of sensitivity, so in this case (Figure 1D & E) the aged *mdx* EDL has become less sensitive to activation by Ca^2+^ as a result of the eccentric contractions consistent with our earlier preliminary findings with the dKO mouse (Hogarth *et al*., 2017). The mouse EDL muscle contains a mixture of fast-twitch fibres ∼79% 2B, ∼16% type 2X and ∼4% type 2A (Hettige *et al*., 2020) and it has been shown that fast-twitch fibres, especially the largest diameter fastest fibre type 2B (Seto *et al*., 2011) are highly susceptible to EC induced force deficits (McCully & Faulkner, 1985; Head *et al*., 1992; Head *et al*., 1994; Chan *et al*., 2007; Schiaffino & Reggiani, 2011; Allen *et al*., 2016; Kiriaev *et al*., 2018), and normally contain α-actinin-3 in the Z-discs (Schiaffino & Reggiani, 2011) which will be the primary transductor of the eccentric force. Given this, it is likely that the absence of α-actinin-3 in the dKO mice has conferred the fast-twitch fibres with some degree of protection from eccentric damage (see discussion below) and hence there is greater damage to the fast-twitch fibres in the aged *mdx* EDL which presents itself as a drop in sensitivity as a result of the remaining force is being generated by the pool of “slower” fast-twitch fibres that remain functional after the eccentric damage. The force frequency parameter values presented here for 12 month old *mdx* EDL (Figure 1) are consistent with the values we reported earlier in aged *mdx* EDL muscles from mice in this age range (Williams *et al*., 1993; Chan *et al*., 2007; Kiriaev *et al*., 2018).

### dKO EDL muscles from 12 month old animals are protected from eccentric contraction induced force deficits despite undergoing a greater “work done” during eccentric contractions

In the dystrophinopathies absence of dystrophin triggers skeletal muscle necrosis followed by regeneration, this process continues throughout the life span of *mdx* mice and DMD boys. The regenerated skeletal muscle fibres are characterised by the presence of centralised nuclei and increasing numbers of abnormal branched fibres, which increase in both number and complexity as the mouse or boy ages. In the later parts of the *mdx* mouse life (12-24 months), some fibres can have up to 10 branches (Kiriaev *et al*., 2018) in a single syncytium with all branched fibres have a single neuromuscular junction (Faber *et al*., 2014; Pratt *et al*., 2015; Massopust *et al*., 2020). In fast-twitch muscles, we and others have found a correlation between the extent and complexity of fibre branching and susceptibility to damage from eccentric contractions (Head *et al*., 1990; Head *et al*., 1992; Chan *et al*., 2007; Head, 2010; Chan *et al*., 2011; Head, 2012; Pichavant *et al*., 2015; Kiriaev *et al*., 2018). In boys with DMD, extensive fibre branching has been associated with an increase in pathological symptoms, such as a reduction in mobility (Bell & Conen, 1968; Schmalbruch, 1984). We have previously provided evidence that in the absence of α-actinin-3 there is a reduced susceptibility to stretch-induced damage in fast-twitch EDL dystrophic muscles from aged 12 month *mdx* mice, however, in young 8 week old mice the absence of α-actinin-3 confers no protection from EC force deficits (Hogarth *et al*., 2017). It is important to note that in these 8 week old muscles 100% of the EDL fibres have undergone at least one round of degeneration and regeneration (Duddy *et al*., 2015), these mice are also from the same α-actinin-3 deficient backgrounds with muscles lacking dystrophin from the inner surface of the sarcolemma.

Here, we extend our earlier finding that absence of α-actinin-3 protects the 12 month fast-twitch *mdx* EDL muscles from eccentric force deficits even when the work done by our EC protocol is markedly larger in the dKO compared to the more severely damaged aged *mdx* muscles (Figure 2C). Work done has been a parameter reported previously when comparing EC damage in dystrophic studies (Stevens & Faulkner, 2000; Lynch *et al*., 2001a; Lynch *et al*., 2001b; Gregorevic *et al*., 2002; Faulkner *et al*., 2008; Kiriaev *et al*., 2018; Lindsay *et al*., 2020). Given that less work was done on aged *mdx* muscles compared with dKO during ECs means it is likely that the protective effect of the absence of α-actinin-3 is even larger than we report here.

### There are less complex branched fibres in EDL muscles from dKO mice which correlates with the protection from eccentric contraction force deficits

In the dystrophinopathies the number and complexity of pathologically branched fibres increases with age (Head *et al*., 1990; Head *et al*., 1992; Chan *et al*., 2007; Lovering *et al*., 2009; Head, 2010; Chan *et al*., 2011; Head, 2012; Buttgereit *et al*., 2013; Faber *et al*., 2014; Pichavant & Pavlath, 2014; Duddy *et al*., 2015; Pichavant *et al*., 2015; Kiriaev *et al*., 2018; Ben Larbi *et al*., 2021). A significant number of published reports have demonstrated a correlation between the extent and complexity of fibre branching in fast-twitch fibres and increased susceptibility to damage from eccentric contractions (Head *et al*., 1992; Dellorusso *et al*., 2001; Chan *et al*., 2007; Pichavant *et al*., 2015; Kiriaev *et al*., 2018). Branching is an important and often overlooked pathological feature of the muscles in the dystrophinopathies and it has been shown that branching is responsible for the hypertrophy of fast-twitch skeletal muscles reported in the *mdx* mouse (Faber *et al*., 2014). We propose the increased susceptibility to injury during eccentric contractions in aged fast-twitch *mdx* muscles is due to morphological changes (branching) of fibres which lead to weakened regions (branch points) that are the site of pathology and acute rupture during contraction (Figure 6), rather than as a direct consequence of the absence of dystrophin (Head, 2010; Chan *et al*., 2011; Head, 2012; Kiriaev *et al*., 2018). In support of this, Pavlaths group have demonstrated that reducing fibre branching in the *mdx* mouse induced by expression of the muscle specific olfactory receptor mOR23 protects against eccentric damage (Pichavant *et al*., 2015). In our initial study (Hogarth *et al*., 2017), we showed that it was only in 12 month dKO mice that the absence of α-actinin-3 was protective against eccentric damage, at younger ages the absence of α-actinin-3 conferred no protection despite the absence of dystrophin in both age groups. Here we show the protective effect of α-actinin-3 in *mdx* fast-twitch muscles at 12 months is even greater than we previously reported (Hogarth *et al*., 2017), furthermore, we show the *Actn3KO* protection is correlated with a reduction in both the number and complexity of branched fibres present in the *mdx* mouse at this age (Figures 3-5).

### Eccentric force loss is gradual and reduced in dKO muscles

Further investigation into the isometric force loss at on each of the three eccentric contractions used in the current study reveal that the force deficit in dKO muscles was graded (Figure 2A). The aged dKO muscles lost ∼35% of their starting force on the first contraction, dramatically different to the ∼80% in aged *mdx* muscles (Figure 2B left). This supports our earlier studies where we showed it was only the aged *mdx* EDL muscles which lost the majority of their force on the first EC (Head *et al*., 1992; Chan *et al*., 2007; Chan *et al*., 2011; Kiriaev *et al*., 2018). We have proposed that this abrupt EC induced force loss in the aged *mdx* mouse is correlated with fibre rupture and tearing at branch points (Kiriaev *et al*., 2018). The gradual force loss observed in dKO muscle resemble the drops in isometric force following EC seen in WT muscles with no branching (Morgan, 1990; Lieber *et al*., 1991; Brooks, 1998; Froehner *et al*., 2014; Kiriaev *et al*., 2018; Kiriaev *et al*., 2021a) and young *mdx* muscles (Kiriaev *et al*., 2018; Kiriaev *et al*., 2021a) which contain a simpler branching morphology, lacking the complex branches observed in aged *mdx* animals that cause the abrupt loss in isometric force.

### Why does the absence of α-actinin-3 reduce fibre branching in the mdx EDL?

Our group has previously reported that *Actn3* polymorphism switches the metabolism in type 2B fibres towards a slower oxidative metabolic state and reduces the 2B fibre diameters compared with WT muscles (Chan *et al*., 2008; Macarthur *et al*., 2008; Seto *et al*., 2011). There would be less shear stress on the fast-twitch fibres in dKO, thus protecting them from EC and potentially providing an explanation for the reduction in fibre branching we report here. Our argument here is that the *Actn3* polymorphism switch of the 2B fibres to a slow-twitch metabolism is protective against EC force deficits, this notion is further supported by our recent publication showing that *mdx* slow-twitch soleus muscles are protected from eccentric force deficits throughout the life span of the *mdx* mouse due to reduced numbers and complexity of branched fibres relative to EDL muscles (Kiriaev *et al*., 2021a). Furthermore, there has been a long history in the muscular dystrophy field which proposes shifting of the fast skeletal muscle fibres to a slow profile will confer protection to DMD boys (for a review see Hardee *et al*. (2021)). Here we provide evidence that *Actn3* polymorphism which switches the anerobic metabolic profile of fast 2B fibres in mice to a slower oxidative pathway without altering the expression of the 2B myosin heavy chain, confers a protection from eccentric force deficits in 12 month *mdx* fast-twitch EDL, and this protection is correlated with a reduction in the number and complexity of branched fibres.

### The molecular mechanism of Actn3KO protection is likely due to an increase in SERCA1

It is well accepted that the absence of dystrophin in the dystrophinopathies, like DMD, triggers skeletal muscle fibre necrosis due to pathological [Ca^2+^]_in_ regulation (Allen *et al*., 2016). Several studies have shown that the skeletal muscle pathology in the dystrophin deficient *mdx* mouse can be ameliorated by overexpressing or pharmacologically activating the SR Ca^2+^ pump, SERCA1 (Goonasekera *et al*., 2011; Mázala *et al*., 2015; Nogami *et al*., 2021). Previously, our group has shown that one consequence of the *Actn3KO* is a marked increase of SERCA1 protein in fast-twitch skeletal muscle (Head *et al*., 2015; Garton *et al*., 2018). In this study we have shown that at 12 months of age dKO mice have a two-fold increase in the SR Ca^2+^ pump SERCA1 compared to age matched dystrophin deficient *mdx* mice (Figure 7). This level of increase is similar to that described by Mázala *et al*. (2015) when using SERCA1 overexpression, and is one likely explanation for the protection conferred by the *Actn3* polymorphism in our dKO mouse. Furthermore, this work provides additional support for the physiological activation of SERCA1 in the skeletal muscle as a potential novel therapeutic target for the treatment of DMD.

### If branches are weak, why do they persist in the dystrophic muscle?

We have hypothesized (Chan *et al*., 2007; Head, 2010; Chan & Head, 2011; Kiriaev *et al*., 2018) that branches are relatively stable within the intact muscle until their number and complexity reach ‘tipping point’. Below this “tipping point” a muscle can contain branched fibres with no loss of mechanical stability (see Kiriaev *et al*. (2021a)). Past “tipping point” fibre branching makes the fibre susceptible to mechanical strains which elicit rupture at one or more branch nodes in the branched fibre syncytium and results in catastrophic failure of the branched fibre unit. If these structural branch failures occur during a strong EC, like the ones we use here *in vitro*, there will be a positive feedback loop placing additional stress on the remaining complex branched fibres and as they rupture so will the remaining branched fibres which have passed ‘tipping point’. However, if the mouse does not encounter eccentric strains of this magnitude these dystrophin-deficient animals can live largely asymptomatically past 108 weeks of age. It is only in the old *mdx* mouse that we start to see them die earlier than age matched controls (Chamberlain *et al*., 2007; Li *et al*., 2009) and we correlate this shortened life span with the degree and complexity of muscle fibre branching passing the “tipping point” where a significant proportion of branches can no longer withstand the normal stresses and strains of muscle contraction.

### Summary

The total absence of α-actinin-3 from the Z-discs of fast-twitch skeletal muscle fibres results in a switch to an oxidative metabolic pathway and increases in the sarcoplasmic reticulum Ca^2+^ pump protein SERCA1. These changes are correlated with a reduction in the occurrence of fast-twitch muscle fibre branching in aged dKO mice, which we propose confers the fast-twitch EDL muscles with some protection from eccentric contraction induced force deficits. These results have implications for disease progression in the ∼16% of boys with dystrophin deficient DMD who also have the common null polymorphism (R577X).

## Supplementary figure

**Supplementary Figure 1.**
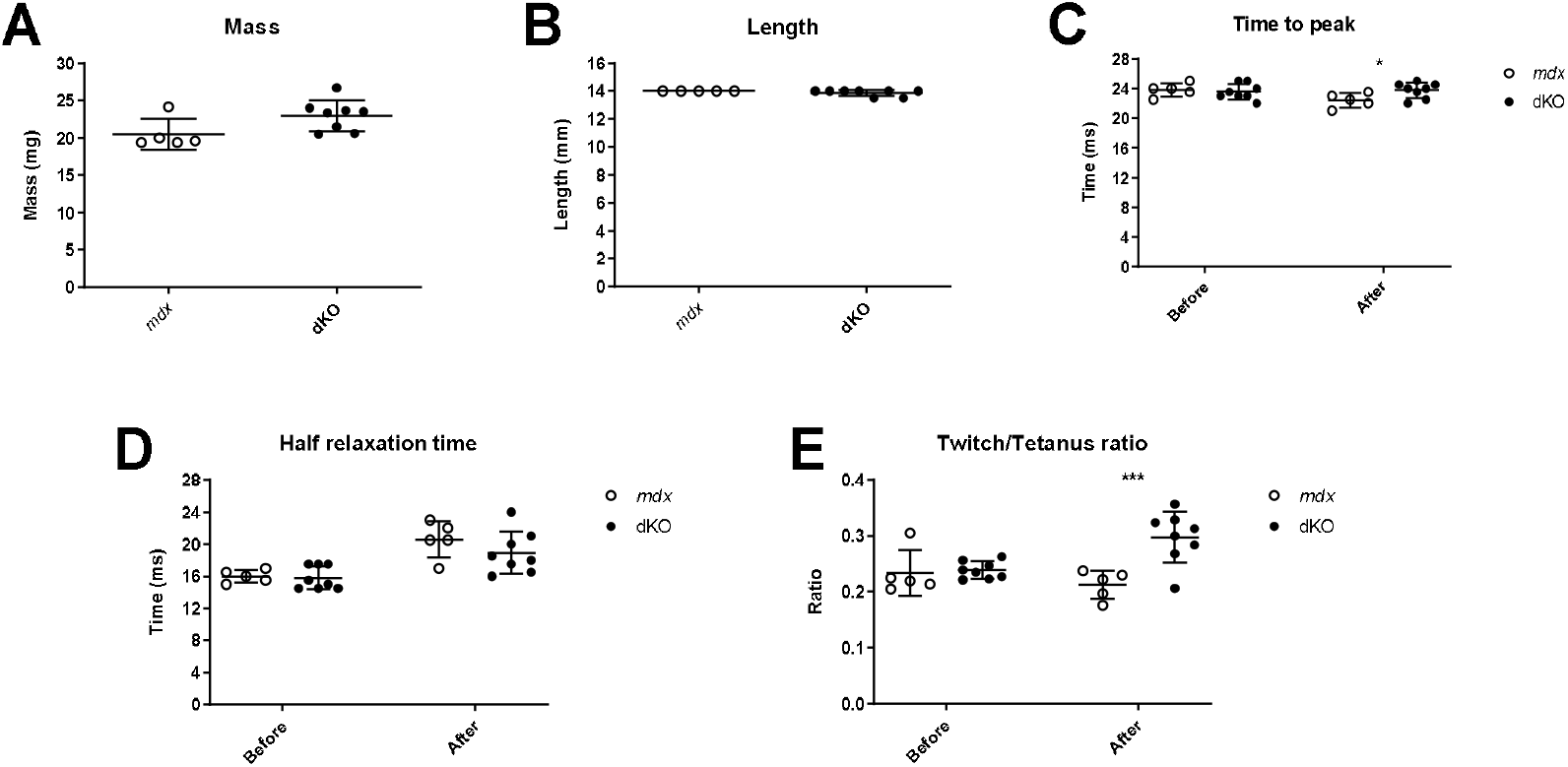
Scatterplots of data presented in Table 1 showing the general physical properties and twitch kinetics of aged *mdx* and dKO EDL muscles. A: Muscle mass, B: Muscle optimal length producing maximal force, C: Time to peak before and after EC, D: Half relaxation time before and after EC and D: twitch to tetanus ratio before and after EC. All data shown are mean ± SD (n=5 Aged *mdx*; n=8 dKO). P-values shown are assessed by unpaired t-test with a significance established at 0.05, ***0.0001<P<0.001 and *0.01<P<0.05.

## References

Aartsma-Rus A & Muntoni F. (2013). 194th ENMC international workshop. 3rd ENMC workshop on exon skipping: Towards clinical application of antisense-mediated exon skipping for Duchenne muscular dystrophy 8–10 December 2012, Naarden, The Netherlands. Neuromuscular Disorders 23, 934–944 DOI.10.1016/j.nmd.2013.06.369

Allen DG, Whitehead NP & Froehner SC. (2016). Absence of Dystrophin Disrupts Skeletal Muscle Signaling: Roles of Ca2+, Reactive Oxygen Species, and Nitric Oxide in the Development of Muscular Dystrophy. Physiol Rev 96, 253–305 DOI.10.1152/physrev.00007.2015

Bell CD & Conen PE. (1968). Histopathological changes in Duchenne muscular dystrophy. Journal of the Neurological Sciences 7, 529–544 DOI.10.1016/0022-510X(68)90058-0

Ben Larbi S, Saclier M, Fessard A, Juban G & Chazaud B. (2021). Histological Analysis of Tibialis Anterior Muscle of DMDmdx4Cv Mice from 1 to 24 Months. J Neuromuscul Dis 8, 513–524 DOI.10.3233/JND-200562

Bengtsson NE, Seto JT, Hall JK, Chamberlain JS & Odom GL. (2016). Progress and prospects of gene therapy clinical trials for the muscular dystrophies. Human Molecular Genetics 25, R9–R17 DOI.10.1093/hmg/ddv420

Berman Y & North KN. (2010). A gene for speed: the emerging role of alpha-actinin-3 in muscle metabolism. Physiology (Bethesda) 25, 250–259 DOI.10.1152/physiol.00008.2010

Brooks SV. (1998). Rapid recovery following contraction-induced injury to in situ skeletal muscles in mdx mice. J Muscle Res Cell Motil 19, 179–187 DOI.10.1023/a:1005364713451

Brooks SV, Zerba E & Faulkner JA. (1995). Injury to muscle fibres after single stretches of passive and maximally stimulated muscles in mice. J Physiol 488 (Pt 2), 459–469 DOI.10.1113/jphysiol.1995.sp020980

Buttgereit A, Weber C, Garbe CS & Friedrich O. (2013). From chaos to split-ups - SHG microscopy reveals a specific remodelling mechanism in ageing dystrophic muscle. The Journal of Pathology 229, 477–485 DOI.10.1002/path.4136

Ceco E & McNally EM. (2013). Modifying muscular dystrophy through transforming growth factor-β. FEBS Journal 280, 4198–4209 DOI.10.1111/febs.12266

Celichowski J & Grottel K. (1993). Twitch/tetanus ratio and its relation to other properties of motor units. Neuroreport 5, 201–204 DOI.10.1097/00001756-199312000-00003

Chamberlain JS, Metzger J, Reyes M, Townsend D & Faulkner JA. (2007). Dystrophin-deficient mdx mice display a reduced life span and are susceptible to spontaneous rhabdomyosarcoma. The FASEB Journal 21, 2195–2204 DOI.10.1096/fj.06-7353com

Chan S & Head SI. (2011). The role of branched fibres in the pathogenesis of Duchenne muscular dystrophy. Exp Physiol 96, 564–571 DOI.10.1113/expphysiol.2010.056713

Chan S, Head SI & Morley JW. (2007). Branched fibers in dystrophic mdx muscle are associated with a loss of force following lengthening contractions. Am J Physiol Cell Physiol 293, C985–992 DOI.10.1152/ajpcell.00128.2007

Chan S, Seto JT, Houweling PJ, Yang N, North KN & Head SI. (2011). Properties of extensor digitorum longus muscle and skinned fibers from adult and aged male and femaleActn3knockout mice. Muscle & Nerve 43, 37–48 DOI.10.1002/mus.21778

Chan S, Seto JT, MacArthur DG, Yang N, North KN & Head SI. (2008). A gene for speed: contractile properties of isolated whole EDL muscle from an alpha-actinin-3 knockout mouse. Am J Physiol Cell Physiol 295, C897–904 DOI.10.1152/ajpcell.00179.2008

Dellorusso C, Crawford RW, Chamberlain JS & Brooks SV. (2001). Tibialis anterior muscles in mdx mice are highly susceptible to contraction-induced injury. Journal of Muscle Research and Cell Motility 22, 467–475 DOI.10.1023/a:1014587918367

Duddy W, Duguez S, Johnston H, Cohen TV, Phadke A, Gordish-Dressman H, Nagaraju K, Gnocchi V, Low S & Partridge T. (2015). Muscular dystrophy in the mdx mouse is a severe myopathy compounded by hypotrophy, hypertrophy and hyperplasia. Skeletal Muscle 5, 16 DOI.10.1186/s13395-015-0041-y

Faber RM, Hall JK, Chamberlain JS & Banks GB. (2014). Myofiber branching rather than myofiber hyperplasia contributes to muscle hypertrophy in mdx mice. Skeletal Muscle 4, 10 DOI.10.1186/2044-5040-4-10

Fairclough RJ, Wood MJ & Davies KE. (2013). Therapy for Duchenne muscular dystrophy: renewed optimism from genetic approaches. Nature Reviews Genetics 14, 373–378 DOI.10.1038/nrg3460

Faulkner JA, Ng R, Davis CS, L. S & Chamberlain JS. (2008). Diaphragm muscle strip preparation for evaluation of gene therapies in mdx mice. Clin Exp Pharmacol Physiol 35, 725–729 DOI.10.1111/j.1440-1681.2007.04865.x

Froehner SC, Reed SM, Anderson KN, Huang PL & Percival JM. (2014). Loss of nNOS inhibits compensatory muscle hypertrophy and exacerbates inflammation and eccentric contraction-induced damage in mdx mice. Human Molecular Genetics 24, 492–505 DOI.10.1093/hmg/ddu469

Garton FC, Houweling PJ, Vukcevic D, Meehan LR, Lee FXZ, Lek M, Roeszler KN, Hogarth MW, Tiong CF, Zannino D, Yang N, Leslie S, Gregorevic P, Head SI, Seto JT & North KN. (2018). The Effect of ACTN3 Gene Doping on Skeletal Muscle Performance. Am J Hum Genet 102, 845–857 DOI.10.1016/j.ajhg.2018.03.009

Goonasekera SA, Lam CK, Millay DP, Sargent MA, Hajjar RJ, Kranias EG & Molkentin JD. (2011). Mitigation of muscular dystrophy in mice by SERCA overexpression in skeletal muscle. Journal of Clinical Investigation 121, 1044–1052 DOI.10.1172/jci43844

Gregorevic P, Plant DR, Leeding KS, Bach LA & Lynch GS. (2002). Improved Contractile Function of the mdx Dystrophic Mouse Diaphragm Muscle after Insulin-Like Growth Factor-I Administration. The American Journal of Pathology 161, 2263–2272 DOI.10.1016/s0002-9440(10)64502-6

Hardee JP, Martins KJB, Miotto PM, Ryall JG, Gehrig SM, Reljic B, Naim T, Chung JD, Trieu J, Swiderski K, Philp AM, Philp A, Watt MJ, Stroud DA, Koopman R, Steinberg GR & Lynch GS. (2021). Metabolic remodeling of dystrophic skeletal muscle reveals biological roles for dystrophin and utrophin in adaptation and plasticity. Molecular Metabolism 45, 101157 DOI.10.1016/j.molmet.2020.101157

Head S, Williams D & Stephenson G. (1994). Increased susceptibility of EDL muscles from mdx mice to damage induced by contraction with stretch. J Muscle Res Cell Motil 15, 490–492

Head SI. (2010). Branched fibres in old dystrophic mdx muscle are associated with mechanical weakening of the sarcolemma, abnormal Ca2+ transients and a breakdown of Ca2+ homeostasis during fatigue. Exp Physiol 95, 641–656 DOI.10.1113/expphysiol.2009.052019

Head SI. (2012). A Two Stage Model of Skeletal Muscle Necrosis in Muscular Dystrophy – The Role of Fiber Branching in the Terminal Stage. In Muscular Dystrophy, ed. Hegde M, pp. 475–498.10.5772/31880.2012

Head SI, Chan S, Houweling PJ, Quinlan KGR, Murphy R, Wagner S, Friedrich O & North KN. (2015). Altered Ca2+ Kinetics Associated with α-Actinin-3 Deficiency May Explain Positive Selection for ACTN3 Null Allele in Human Evolution. PLOS Genetics 11, e1004862 DOI.10.1371/journal.pgen.1004862

Head SI, Stephenson DG & Williams DA. (1990). Properties of enzymatically isolated skeletal fibres from mice with muscular dystrophy. J Physiol 422, 351–367 DOI.10.1113/jphysiol.1990.sp017988

Head SI, Williams DA & Stephenson DG. (1992). Abnormalities in structure and function of limb skeletal muscle fibres of dystrophic mdx mice. Proc Biol Sci 248, 163–169 DOI.10.1098/rspb.1992.0058

Hettige P, Tahir U, Nishikawa KC & Gage MJ. (2020). Comparative analysis of the transcriptomes of EDL, psoas, and soleus muscles from mice. BMC Genomics 21, 808 DOI.10.1186/s12864-020-07225-2

Hoffman EP & Connor EM. (2013). Orphan drug development in muscular dystrophy: update on two large clinical trials of dystrophin rescue therapies. Discov Med 16, 233–239

Hoffman EP & McNally EM. (2014). Exon-skipping therapy: a roadblock, detour, or bump in the road? Sci Transl Med 6, 230fs214 DOI.10.1126/scitranslmed.3008873

Hogarth MW, Houweling PJ, Thomas KC, Gordish-Dressman H, Bello L, Pegoraro E, Hoffman EP, Head SI & North KN. (2017). Evidence for ACTN3 as a genetic modifier of Duchenne muscular dystrophy. Nature Communications 8, 14143 DOI.10.1038/ncomms14143

Houweling PJ, Papadimitriou ID, Seto JT, Pérez LM, Coso JD, North KN, Lucia A & Eynon N. (2018). Is evolutionary loss our gain? The role of ACTN3 p.Arg577Ter (R577X) genotype in athletic performance, ageing, and disease. Human Mutation 39, 1774–1787 DOI.10.1002/humu.23663

Hunter KD & Faulkner JA. (1997). Pliometric contraction-induced injury of mouse skeletal muscle: effect of initial length. J Appl Physiol 82, 278–283 DOI.10.1152/jappl.1997.82.1.278

Kiriaev L, Kueh S, Morley JW, Houweling PJ, Chan S, North KN & Head SI. (2021a). Dystrophin-negative slow-twitch soleus muscles are not susceptible to eccentric contraction induced injury over the lifespan of the mdx mouse. Am J Physiol Cell Physiol 0, null DOI.10.1152/ajpcell.00122.2021

Kiriaev L, Kueh S, Morley JW, North KN, Houweling PJ & Head SI. (2018). Branched fibers from old fast-twitch dystrophic muscles are the sites of terminal damage in muscular dystrophy. Am J Physiol Cell Physiol 314, C662–C674 DOI.10.1152/ajpcell.00161.2017

Kiriaev L, Kueh S, Morley JW, North KN, Houweling PJ & Head SI. (2021b). Lifespan analysis of dystrophic mdx fast-twitch muscle morphology and its impact on contractile function. Western Sydney University.’preprint 10.1101/2021.09.07.459226

Lee FXZ, Houweling PJ, North KN & Quinlan KGR. (2016). How does α-actinin-3 deficiency alter muscle function? Mechanistic insights into ACTN3, the ‘gene for speed’. Biochimica et Biophysica Acta (BBA) - Molecular Cell Research 1863, 686–693 DOI.10.1016/j.bbamcr.2016.01.013

Li D, Long C, Yue Y & Duan D. (2009). Sub-physiological sarcoglycan expression contributes to compensatory muscle protection in mdx mice. Human Molecular Genetics 18, 1209–1220 DOI.10.1093/hmg/ddp015

Lieber RL, Woodburn TM & Friden J. (1991). Muscle damage induced by eccentric contractions of 25% strain. Journal of Applied Physiology 70, 2498–2507 DOI.10.1152/jappl.1991.70.6.2498

Lindsay A, Baumann CW, Rebbeck RT, Yuen SL, Southern WM, Hodges JS, Cornea RL, Thomas DD, Ervasti JM & Lowe DA. (2020). Mechanical factors tune the sensitivity of mdx muscle to eccentric strength loss and its protection by antioxidant and calcium modulators. Skeletal Muscle 10, 3 DOI.10.1186/s13395-020-0221-2

Lovering RM, Michaelson L & Ward CW. (2009). Malformed mdx myofibers have normal cytoskeletal architecture yet altered EC coupling and stress-induced Ca2+ signaling. American Journal of Physiology-Cell Physiology 297, C571–C580 DOI.10.1152/ajpcell.00087.2009

Lu QL, Cirak S & Partridge T. (2014). What Can We Learn From Clinical Trials of Exon Skipping for DMD? Mol Ther Nucleic Acids 3, e152 DOI.10.1038/mtna.2014.6

Lynch GS, Hinkle RT, Chamberlain JS, Brooks SV & Faulkner JA. (2001a). Force and power output of fast and slow skeletal muscles from mdx mice 6-28 months old. J Physiol 535, 591–600 DOI.10.1111/j.1469-7793.2001.00591.x

Lynch GS, Hinkle RT & Faulkner JA. (2001b). Force and power output of diaphragm muscle strips from mdx and control mice after clenbuterol treatment. Neuromuscular Disorders 11, 192–196 DOI.10.1016/s0960-8966(00)00170-x

Macarthur DG, Seto JT, Chan S, Quinlan KGR, Raftery JM, Turner N, Nicholson MD, Kee AJ, Hardeman EC, Gunning PW, Cooney GJ, Head SI, Yang N & North KN. (2008). An Actn3 knockout mouse provides mechanistic insights into the association between -actinin-3 deficiency and human athletic performance. Human Molecular Genetics 17, 1076–1086 DOI.10.1093/hmg/ddm380

Macarthur DG, Seto JT, Raftery JM, Quinlan KG, Huttley GA, Hook JW, Lemckert FA, Kee AJ, Edwards MR, Berman Y, Hardeman EC, Gunning PW, Easteal S, Yang N & North KN. (2007). Loss of ACTN3 gene function alters mouse muscle metabolism and shows evidence of positive selection in humans. Nature Genetics 39, 1261–1265 DOI.10.1038/ng2122

Massopust RT, Lee YI, Pritchard AL, Nguyen VM, McCreedy DA & Thompson WJ. (2020). Lifetime analysis of mdx skeletal muscle reveals a progressive pathology that leads to myofiber loss. Sci Rep 10, 17248 DOI.10.1038/s41598-020-74192-9

Mázala DAG, Pratt SJP, Chen D, Molkentin JD, Lovering RM & Chin ER. (2015). SERCA1 overexpression minimizes skeletal muscle damage in dystrophic mouse models. American Journal of Physiology-Cell Physiology 308, C699–C709 DOI.10.1152/ajpcell.00341.2014

McCully KK & Faulkner JA. (1985). Injury to skeletal muscle fibers of mice following lengthening contractions. J Appl Physiol (1985) 59, 119–126 DOI.10.1152/jappl.1985.59.1.119

Morgan DL. (1990). New insights into the behavior of muscle during active lengthening. Biophysical journal 57, 209–221 DOI.10.1016/S0006-3495(90)82524-8

Nogami KI, Maruyama Y, Sakai-Takemura F, Motohashi N, Elhussieny A, Imamura M, Miyashita S, Ogawa M, Noguchi S, Tamura Y, Kira J-I, Aoki Y, Takeda SI & Miyagoe-Suzuki Y. (2021). Pharmacological activation of SERCA ameliorates dystrophic phenotypes in dystrophin-deficient mdx mice. Human Molecular Genetics 30, 1006–1019 DOI.10.1093/hmg/ddab100

North KN, Yang N, Wattanasirichaigoon D, Mills M, Easteal S & Beggs AH. (1999). A common nonsense mutation results in α-actinin-3 deficiency in the general population. Nature Genetics 21, 353–354 DOI.10.1038/7675

Patel TJ & Lieber RL. (1997). Force transmission in skeletal muscle: from actomyosin to external tendons. Exerc Sport Sci Rev 25, 321–363

Peczkowski KK, Rastogi N, Lowe J, Floyd KT, Schultz EJ, Karaze T, Davis JP, Rafael-Fortney JA & Janssen PML. (2020). Muscle Twitch Kinetics Are Dependent on Muscle Group, Disease State, and Age in Duchenne Muscular Dystrophy Mouse Models. Front Physiol 11, 568909 DOI.10.3389/fphys.2020.568909

Pichavant C, Burkholder TJ & Pavlath GK. (2015). Decrease of myofiber branching via muscle-specific expression of the olfactory receptor mOR23 in dystrophic muscle leads to protection against mechanical stress. Skelet Muscle 6, 2 DOI.10.1186/s13395-016-0077-7

Pichavant C & Pavlath GK. (2014). Incidence and severity of myofiber branching with regeneration and aging. Skeletal Muscle 4, 9 DOI.10.1186/2044-5040-4-9

Pratt SJP, Valencia AP, L. GK, Shah SB & Lovering RM. (2015). Pre- and postsynaptic changes in the neuromuscular junction in dystrophic mice. Front Physiol 6, 252–252 DOI.10.3389/fphys.2015.00252

Quinlan KGR, Seto JT, Turner N, Vandebrouck A, Floetenmeyer M, Macarthur DG, Raftery JM, Lek M, Yang N, Parton RG, Cooney GJ & North KN. (2010). α-Actinin-3 deficiency results in reduced glycogen phosphorylase activity and altered calcium handling in skeletal muscle. Human Molecular Genetics 19, 1335–1346 DOI.10.1093/hmg/ddq010

Rodino-Klapac LR, Mendell JR & Sahenk Z. (2013). Update on the Treatment of Duchenne Muscular Dystrophy. Current Neurology and Neuroscience Reports 13, 332 DOI.10.1007/s11910-012-0332-1

Schiaffino S & Reggiani C. (2011). Fiber types in mammalian skeletal muscles. Physiol Rev 91, 1447–1531 DOI.10.1152/physrev.00031.2010

Schmalbruch H. (1984). Regenerated muscle fibers in Duchenne muscular dystrophy: a serial section study. Neurology 34, 60–65 DOI.10.1212/wnl.34.1.60

Seto JT, Lek M, Quinlan KGR, Houweling PJ, Zheng XF, Garton F, Macarthur DG, Raftery JM, Garvey SM, Hauser MA, Yang N, Head SI & North KN. (2011). Deficiency of α-actinin-3 is associated with increased susceptibility to contraction-induced damage and skeletal muscle remodeling. Human Molecular Genetics 20, 2914–2927 DOI.10.1093/hmg/ddr196

Shieh PB. (2015). Duchenne muscular dystrophy: clinical trials and emerging tribulations. Curr Opin Neurol 28, 542–546 DOI.10.1097/WCO.0000000000000243

Stevens ED & Faulkner JA. (2000). The capacity of mdx mouse diaphragm muscle to do oscillatory work. The Journal of Physiology 522, 457–466 DOI.10.1111/j.1469-7793.2000.t01-3-00457.x

Williams DA, Head SI, Lynch GS & Stephenson DG. (1993). Contractile properties of skinned muscle fibres from young and adult normal and dystrophic (mdx) mice. The Journal of Physiology 460, 51–67 DOI.10.1113/jphysiol.1993.sp019458

Wyckelsma VL, Venckunas T, Houweling PJ, Schlittler M, Lauschke VM, Tiong CF, Wood HD, Ivarsson N, Paulauskas H, Eimantas N, Andersson DC, North KN, Brazaitis M & Westerblad. (2021). Loss of α-actinin-3 during human evolution provides superior cold resilience and muscle heat generation. American journal of human genetics 108, 446–457 DOI.10.1016/j.ajhg.2021.01.013

Yang N, MacArthur DG, Gulbin JP, Hahn AG, Beggs AH, Easteal S & North K. (2003). ACTN3 genotype is associated with human elite athletic performance. Am J Hum Genet 73, 627–631 DOI.10.1086/377590

